# Eliminating Malaria Vectors with Precision Guided Sterile Males

**DOI:** 10.1101/2023.07.20.549947

**Authors:** Andrea L. Smidler, Reema A. Apte, James J. Pai, Martha L. Chow, Sanle Chen, Agastya Mondal, Héctor M. Sánchez C., Igor Antoshechkin, John M. Marshall, Omar S. Akbari

## Abstract

Controlling the principal African malaria vector, the mosquito *Anopheles gambiae*, is considered essential to curtail malaria transmission. However existing vector control technologies rely on insecticides, which are becoming increasingly ineffective. Sterile insect technique (SIT) is a powerful suppression approach that has successfully eradicated a number of insect pests, yet the *A. gambiae* toolkit lacks the requisite technologies for its implementation. SIT relies on iterative mass-releases of non-biting, non-driving, sterile males which seek out and mate with monandrous wild females. Once mated, females are permanently sterilized due to mating-induced refractoriness, which results in population suppression of the subsequent generation. However, sterilization by traditional methods renders males unfit, making the creation of precise genetic sterilization methods imperative. Here we develop precision guided Sterile Insect Technique (pgSIT) in the mosquito *A. gambiae* for inducible, programmed male-sterilization and female-elimination for wide scale use in SIT campaigns. Using a binary CRISPR strategy, we cross separate engineered Cas9 and gRNA strains to disrupt male-fertility and female-essential genes, yielding >99.5% male-sterility and >99.9% female-lethality in hybrid progeny. We demonstrate that these genetically sterilized males have good longevity, are able to induce population suppression in cage trials, and are predicted to eliminate wild *A. gambiae* populations using mathematical models, making them ideal candidates for release. This work provides a valuable addition to the malaria genetic biocontrol toolkit, for the first time enabling scalable SIT-like confinable suppression in the species.

## Introduction

Malaria is a deadly parasitic disease that kills a child every minute (1), and is the root cause and consequence of poverty in Africa (2). While widespread vaccine distribution recently began to avert the worst disease outcomes (3, 4), eradication remains elusive. Controlling the principal African malaria mosquito vector, *Anopheles gambiae*, promises to facilitate control and perhaps even elimination of disease transmission in the most highly infected areas. However, currently implemented control methods including insecticide-based technologies, and environmental controls, are increasingly less effective with the list of resistant populations growing yearly (5). Therefore novel non-insecticidal control measures are needed to curb the spread of disease.

To fill this critical niche, genetic vector control technologies are being developed in *Anopheles gambiae*. In this species, the technology farthest down the developmental pipeline are gene drives – selfish genetic elements capable of unilaterally engineering entire wild populations (6–8). However, due to their self-autonomous spread (9), and propensity for breakage via unavoidable generation of resistant alleles (10, 11), gene drives have unsurprisingly spurred scientific, social, ethical, economic, ecological, practical, and political concerns hindering their potential roll-out (12–17). To expedite approval and save lives, and to provide more durable, and controllable, immediate options, it is imperative we develop alternative vector control measures that have safe track records. Sterile Insect Technique (SIT) is one such potential technology, as it has been used to eliminate the tsetse fly, screwworm, melon fly, medfly, and *Aedes* pest populations to great effect (18–23). Requiring both a male-sterilization and female-removal component, SIT acts through the mass-releases of infertile males which naturally locate, copulate with, and sterilize their monandrous female mates. Because the control agent is an insect rather than a traditional pesticide, it has minimal off-target effects. Furthermore, SIT males can seek out and sterilize females in cryptic locations that insecticides may miss, and are the sex that does not drink blood nor spread disease. As a result, SIT acts as a species-specific and chemical-free ‘organic’ insecticide that has potential to enable automated, safe, scalable, and controllable suppression when adapted to *A. gambiae*.

Building a scalable genetic SIT system in anophelines requires creating and combining precise male-sterilization and female-elimination systems, a process not yet successfully undertaken in the species. Sterilization by traditional chemo- or radio-sterilization methods unfortunately impairs male fitness (24–29). Oxidation of somatic DNA, lipids, and proteins (30), causes reduced emergence (18, 27), longevity (25, 31), sperm production (32), and ability to compete during copulation (25, 26, 33, 34) - a lekking based process where competition is fierce (35). Only partially-sterilizing radiation doses generate sufficiently competitive males in trials, although with compromised population suppression efficacy (36, 37). For these reasons researchers have sought to develop male-sterilization technologies using more precise genetic methods. For example, *A. gambiae* lines have been developed which shred the embryonic X-chromosome (38, 39), or express pro-apoptotic factors in the testis (40), resulting in sterility or offspring killing. However, these lines are difficult to rear in mass because they lack the ability to induce, or repress, the sterility phenotype. In a step towards a more scalable technology, a binary CRISPR system was recently demonstrated in *A. gambiae* which could generate spermless males in a more inducible manner (41). However it caused incomplete genetic sterilization (95%) and lacked a sex-sorting component - a hindrance shared by all *A. gambiae* sterilization technologies to date.

Efforts to develop efficient genetic sexing systems (GSSs) in *A. gambiae* have been fruitful but limited. Historically, to achieve male-only lines scientists employed a genetic sexing strain (42) reliant on Y chromosome-linked resistance to dieldrin. However many of these lines are now extinct (43), and use of dieldrin banned as it is highly neurotoxic (44, 45), impeding implementation. Therefore safe genetic sex-separation systems in *A. gambiae* have thus far been limited to transgenic expression of sex-specific fluorophores (46–48) followed by fluorescent sorting of neonate larvae via COPAS (49). However these fluorescence-based technologies require sorting of the released generation directly prior to release, making an egg-based distribution modality impossible, a desirable feature if scaling to cover vast distances. Systems which shred the X-chromosome have also been generated which yield highly male-biased populations. However these lines unfortunately lack inducibility, or repressibility, making them exceedingly difficult to scale (29, 50). Fortunately, the genetic sexing system Ifegenia (Inherited Female Elimination by Genetically Encoded Nucleases to Interrupt Alleles) was recently developed, which permits egg distribution due to automatic genetic sexing. Ifegenia remarkably kills >99.9% females using a binary CRISPR system to target the female-essential gene *femaleless* (*fle*), producing a robust and inducible GSS through genetic crosses (51). Taken together, there remains high demand for a scalable SIT system that encompasses both highly-penetrant male sterilization and female elimination.

One complete genetic SIT system which combines female elimination and male sterilization is termed precision guided Sterile Insect Technique (pgSIT). It has been successfully developed in *Aedes* and *Drosophila* (52–55), but not yet in an anopheline species. PgSIT induces male sterilization and female elimination in the offspring of a cross between separate Cas9 and gRNA lines that target male-fertility and female-essential genes during development, resulting in an ‘inducible’ system suited to large-scale releases. In this work, we develop a pgSIT system in *Anopheles gambiae* that is highly efficient at sterilizing males and eliminating females. We develop a multi-gRNA line targeting the well-characterized female-essential locus, *doublesexF* (*dsxF*)(6), and male-fertility genes *Zero population growth* (*zpg*)(56) and β*2-tubulin* (β*2*)(48). We demonstrate that crossing this gRNA line to Cas9 yields female-androgenization and robust male-sterilization in the resulting hybrid F1 offspring. We then improve female-elimination by introducing the new GSS, Ifegenia (51), which targets the female-essential gene, *fle,* enabling an egg-based distribution modality. In this more complete pgSIT system, we demonstrate complete female killing (>99.9%), near complete male sterility (>99.5%), efficient population suppression in cage trials, and model-predicted elimination of *A. gambiae* populations in the wild, demonstrating proof-of-principle that pgSIT is a candidate system for confinable vector control of the deadly *A. gambiae* malaria vector.

## Results

### Founding and characterizing gZBD: females are incompletely androgenized, males are highly sterile

To develop a pgSIT system in *A. gambiae*, we built a gRNA-expressing transgene, gZBD, that encodes a *Actin5c-m2Turquoise* marker and five gRNAs: one gRNA targeting the germline-essential gap-junction gene ***z****pg* (*gRNA^zpg.1^*)(41), two gRNAs targeting the sperm motility gene β*2-tubulin* (gRNA^β2.1^,gRNA^β2.2^), and two gRNAs targeting the female-differentiation gene ***d****sxF* (gRNA^dsxF.1^, gRNA^dsxF.2^) (**Figure 1A**). We established two distinct gZBD families, gZBD^A18^ and gZBD^D15^, with different transgene insertion sites and expression profiles, and identified their chromosomal insertion positions by inverse PCR at (chr3L:34188038) and (chr3L:828896) respectively (57). For the Cas9 line, we utilized the Vasa2-Cas9 line (58), VZC, (henceforth referred to as Cas9) characterized by a *3xP3-DsRed* selectable marker. It was selected due to its robust germline expression profile and ability to deposit Cas9 in the embryo, which promotes desired F1 mosaic mutagenesis during development resulting in a phenomenon we coined lethal mosaicism (52).

**Figure 1:**
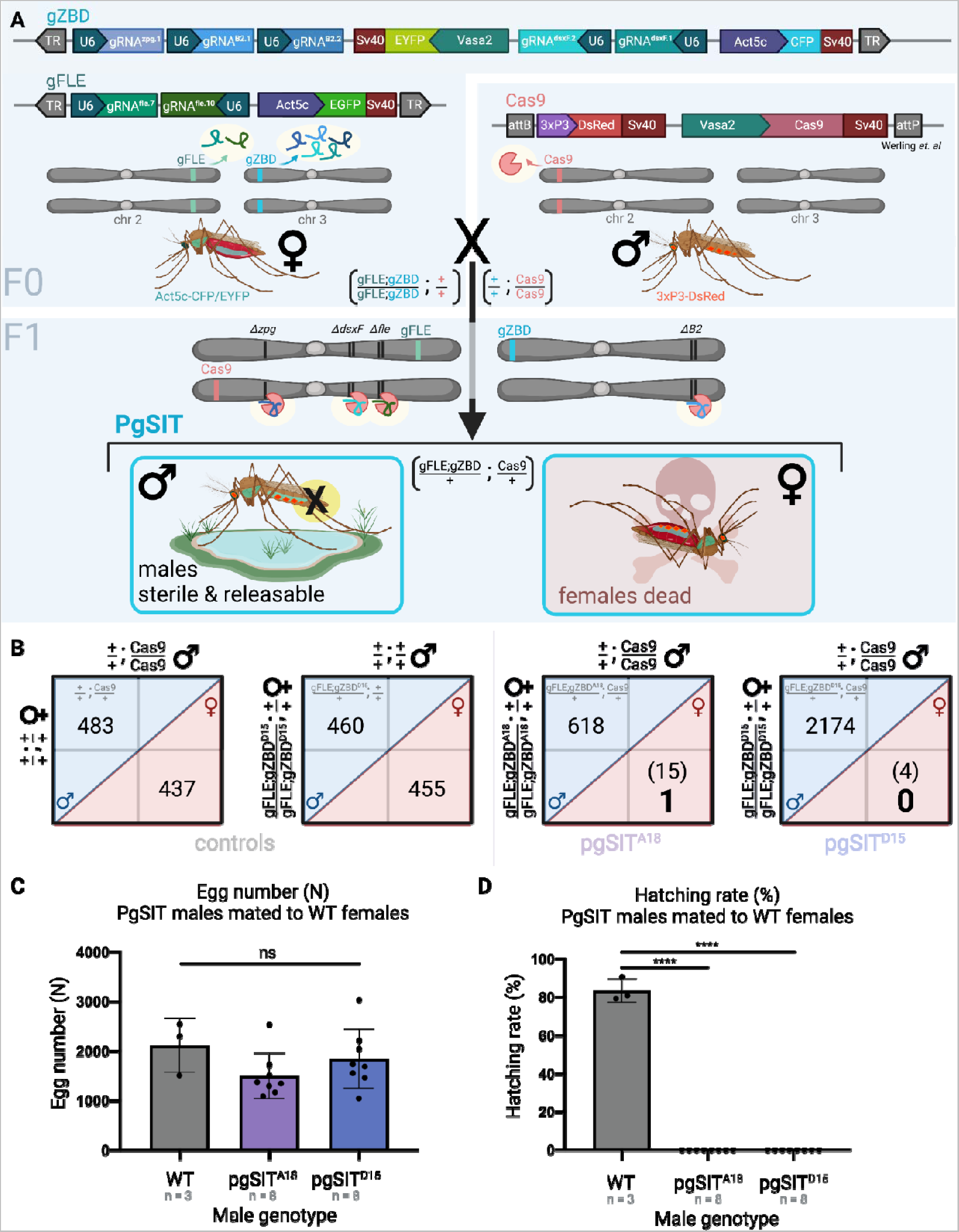
Homozygous pgSIT gRNA females crossed to Cas9 males produce nearly exclusively sterile male F1 offspring. **A)** gZBD transgenics express one gRNA targeting *zero population growth* (*zpg*) (lavender), two gRNAs targeting _J*2-tubulin (*periwinkle), and two gRNAs targeting the female specific exon 5 of *doublesex* (*dsxF*) (teal) under the expression of individual PolIII U6 promoters. Also included is a whole-body fluorescent selectable marker, *Actin5c-m2turquoise* (denoted CFP for brevity), as well as a *Vasa2-EYFP* marker to aid in germline visualization, a marker which was not visible in these lines. gZBD transgenic lines were individually crossed to a second line, gFLE, to generate double homozygous transgenic lines termed (gFLE;gZBD). gFLE targets *femaleless* (*fle*) via two gRNAs also under the expression of the polIII U6 promoter, and includes an *Actin5c-EGFP* cassette for selection by whole-body fluorescence. A third line, Cas9, expresses Cas9 in the germline under the *Vasa2* promoter and includes a *3xP3-DsRed* cassette for selection by central nervous system fluorescence. Crossing (gFLE;gZBD) females to Cas9 males yields trans-heterozygous pgSIT individuals who bear all three transgenes, resulting in active mosaic mutagenesis, and causing female killing and male sterilization. **B)** Among control and pgSIT test crosses, the female-killing phenotype was quantified in the F1 generation, reported as male and female sibling pupa counts. Male and female counts are delineated within blue and red diagonal areas respectively. Control crosses of Cas9 or (gFLE;gZBD^D15^) homozygotes to wild type result in approximately equal F1 male and female pupa counts. Crosses between (gFLE;gZBD^A18^) or (gFLE;gZBD^D15^) homozygous females and Cas9 homozygous males results in significantly reduced F1 female pupa numbers (in parentheses) (p < 0.0001 for both groups, Binomial test). The number of pupae which survived to adulthood to fly are denoted in large bold font. **C)** Crossing 50 pgSIT males to 50 wild type females results in statistically the same number of eggs being laid. Three cage replicates and eight cage replicates shown for wild type control and pgSIT test genotypes respectively. Raw egg counts shown (ns, One Way ANOVA, Dunnetts multiple comparisons test). Mean and SD shown. **D)** Crossing 50 pgSIT males to 50 wild type females results in complete sterilization of females when assayed by hatching rate (n% = n 1 day old larvae /n eggs laid), with high significance compared to the wild type control group (p < 0.0001 for each group, One Way ANOVA, Dunnetts multiple comparisons test). Three cage replicates and eight cage replicates shown for wild type control and pgSIT test genotypes respectively.

We hypothesized that crossing the gZBD and Cas9 lines would yield (+/gZBD; +/Cas9) F1 hybrid offspring with the desired female androgenization and male sterilization mosaic knockout phenotypes. Among the hybrid F1 offspring we identified some intersex (+/gZBD; +/Cas9) females with male claspers on female genitalia indicative of *dsxF* mutagenesis (**Figure S1**)(6). When assaying gZBD^A18^ and gZBD^D15^ individually, we observed only 77% and 68% intersex phenotype penetrance respectively among F1 hybrid offspring (**Table S1**), with some females able to initiate blood feeding (n = 3/26). While this assay involved examining genital claspers and not internal reproductive morphology, it still indicates incomplete androgenization making them unacceptable for vector control.

To determine if (+/gZBD; +/Cas9) males are sterile, we performed crosses of 50 wild type females to 50 F1 (+/gZBD^A18^; +/Cas9) or (+/gZBD^D15^; +/Cas9) males and assayed the hatching rates of their F2 offspring. We observed sterilization of all females mated to (+/gZBD^A18^; +/Cas9) males, with a hatch rate of 0%, and most females mated to (+/gZBD^D15^; +/Cas9) males with an F2 offspring hatch rate of 5.1%, compared to 82.3% hatching rate in wild type controls (**Figure S2, Table S2**). Hatched F2 larvae were verified to express the transgenic fluorescence ratios indicative of (+/gZBD; +/Cas9) paternity, verifying the presence of an ‘escapee’ fertile male. Sequencing these F2 larvae revealed no mutations at the target sites, suggesting incomplete germline mutagenesis in their (+/gZBD; +/Cas9) father. Cumulatively, our data shows that the preliminary pgSIT design robustly sterilizes males with efficiency dependent upon genomic insertion site, but fails to sufficiently incapacitate females.

### Improving female-killing by combining with gFLE

We hypothesized that we could improve the female elimination properties of our system by additionally targeting the recently-discovered female-essential gene *fle* through the introgression of the Ifegenia GSS gRNA line (51). To do this, we separately crossed both gZBD lines into the previously-published gFLE_G_ transgenic line (hereon shortened to gFLE) to produce the doubly homozygous gRNA lines (gFLE;gZBD^A18^) and (gFLE;gZBD^D15^). The gFLE line expresses two gRNAs targeting *fle* and an *Actin5c-EGFP* selectable marker (51) (**Figure 1A**), making it distinguishable from the *Actin5c-m2Turquoise* on gZBD. We previously showed that crossing gFLE males to Cas9 females results in complete female death in the F1 transheterozygous offspring before the pupal stage. Therefore, we hypothesized that (gFLE;gZBD) crossed to Cas9 would produce a robust pgSIT system with the male-sterilizing properties of gZBD and the female-killing properties of gFLE.

### pgSIT (+/gFLE;gZBD; +/Cas9) individuals are mosaic mutants, but some crosses are lethal

Prior pgSIT and Ifegenia systems generated F1 hybrids using F0 Cas9 females (as opposed to males) because they are capable of maternal deposition of Cas9 into F1 embryos, resulting in stronger mosaic mutagenesis and more penetrant phenotypes. For initial verification of mutagenesis we crossed homozygous (gFLE;gZBD) males to homozygous Cas9 females and confirmed mutations in *zpg*, β*2-tubulin*, *dsxF,* and *fle* in F1 embryos (**Figure S3**). However, we observed these crosses resulted in severe F1 mortality at the early larval stage, even though separate F0 crosses between gFLE or gZBD males to Cas9 females were viable. Fortunately, the reciprocal F0 cross using Cas9 males generated viable F1 offspring, and was used to generate the (+/gFLE;gZBD; +/Cas9) genotype used in all subsequent experiments. For simplicity, the hybrid F1 (+/gFLE;gZBD; +/Cas9) genotype is henceforth abbreviated to ‘pgSIT’ genotype, with (+/gFLE;gZBD^A18^; +/Cas9) and (+/gFLE;gZBD^D15^; +/Cas9) shortened to pgSIT^A18^ and pgSIT^D15^ respectively.

### The pgSIT system induces robust female elimination and produces fit sterile males

We next determined if this new pgSIT system is capable of robustly eliminating females and sterilizing males. We observed almost exclusively male pupae among both pgSIT^A18^ and pgSIT^D15^ individuals, indicating robust female elimination (**Figure 1B**). Specifically, for 618 pgSIT^A18^ male pupae scored, 15 female sibling pupae were identified, out of which only one survived to fly; and for 2174 pgSIT^D15^ male pupae scored, 4 female pupae siblings were identified, out of which none survived to fly (**Figure 1B, Table S3**). Consistent with prior work (51), both pgSIT^A18^ and pgSIT^D15^ lines exhibited robust female elimination, 99.8% and 100% and respectively, sufficient to be candidates for field releases. To determine if pgSIT males have high sterility, we crossed 50 pgSIT^A18^ or pgSIT^D15^ males to 50 wild type females and calculated percent fertility of the resulting broods. Out of 16 total cages assayed (800 males total, 400 for each family), zero larvae hatched, demonstrating 100% sterility of both pgSIT^A18^ and pgSIT^D15^ males in these assays (**Figure 1C,D, Table S4**). A more accurate sterility measurement for the population as a whole is >99.5% for each line assuming half of the males were represented in the assay.

Moving forward, we selected pgSIT^D15^ for further characterization, crosses, and analysis due to its strong female-killing and male sterility phenotypes, as well as husbandry considerations. To verify the sterility phenotype, we performed dissections on male pgSIT^D15^ lower reproductive tracts, which revealed the absence of normal testicular tissue (**Figure S4A**). As expected, we observed atrophied testes within (+/gZBD; +/Cas9) controls due to *zpg* and *B2-tubulin* targeting. However, we also observed atrophied and occasionally absent testes among (+/gFLE; +/Cas9) controls, a phenotype not noticed in prior work due to the fertility of the (+/gFLE; +/Cas9) male population as a whole (51). This suggests that targeting all of these genes together may have an additive, or synergistic effect, causing the complete sterility observed in (**Figure 1C**), as opposed to the partial fertility observed in (**Figure S2C**). Taken together, these results demonstrate that pgSIT^D15^ males are sufficiently sterilized to be candidates for SIT by most measures.

For the most effective population suppression, males must be able to mate with and induce mating refractoriness in females, in addition to having high fitness. In *A. gambiae,* refractoriness is induced following the transfer of a gelatinous mating plug to the female during copulation, a structure originally produced by the Male Accessory Glands (MAGs) (59, 60). Dissections revealed that despite lacking testes, pgSIT^D15^ males still developed other important tissues for reproduction such as claspers, an aedegus, and MAGs (**Figure S4A)**. In line with the development of MAGs, we confirmed that pgSIT^D15^ males were able to successfully transfer a mating plug during copulation (**Figure S4B**), indicating females should be refractory to further mating (60, 61). To quantify general pgSIT male fitness, we determined their longevity through survival curve assays (**Figure S4C, Table S5**). These revealed that pgSIT^D15^ males have approximately the same longevity as wild type males (p = ns Mantel Cox test), living slightly but not significantly longer than controls. In summary, these results suggest that pgSIT^D15^ males do not have significant fitness costs that could curtail their longevity and develop all structures critical for reproduction, suggesting they have high fitness overall.

To further characterize the pgSIT^D15^ line, we performed Nanopore sequencing on pooled pgSIT^D15^ males and confirmed the single insertion site of gZBD^D15^ to be within a noncoding region of chr. 3L (NT_078267.5:4828892-4828896). We also validated the previously characterized insertions of gFLE and Cas9 within 2R and 2L, respectively (51, 58). Interestingly, nanopore also showed a large deletion over gRNA^dsxF.1^ within gZBD^D15^ encompassing the U6 promoter and gRNA coding sequence. Sanger sequencing of both gZBD^D15^ and gFLE;gZBD^D15^ individuals confirmed the deletion over gRNA^dsxF.1^ and confirmed that all other gRNA coding sequences on the transgene were intact (**Figure S5**). In line with this, we observed CRISPR-induced mutations in pgSIT^D15^ samples at each gRNA genomic target site except for gRNA^dsx.F1^ (**Figure S3**). While it is unknown exactly when the deletion took place, clear intersex phenotypes were observed during initial line characterization (**Figure S1, Table S1**), but were not observed in the (+/gZBD^D15^; +/Cas9) female siblings of the males used in **Figure S2B**, and may explain the partial phenotypes observed during initial line characterization. This suggests that gRNA^dsxF.1^ had been lost in the interim and that cleavage by gRNA^dsxF.2^ does not cause intersex phenotypes. Regardless, pgSIT^D15^ is still a viable candidate because the fully-penetrant female-killing phenotype caused by *fle*-targeting makes *dsxF*-targeting redundant, rendering gRNA^dsxF.1^ irrelevant to pgSIT system function.

### PgSIT can induce population suppression

We next set out to determine if pgSIT^D15^ males could cause population suppression in cage trials. For this we established competition cages of pgSIT^D15^ males against 50 wild type males at 0:1, 1:1, 2:1, 5:1 or 10:1 ratios, and added 50 wild type females as potential mates. The resulting broods were assayed for percent fertility (**Figure 2A**). Consistent with release ratios required to suppress *Aedes* populations (62), we observed significant population suppression from the 10:1 and 5:1 release ratios (17.4% and 29.7% mean hatching rate, both p < 0.0001), and non-significant, less pronounced, suppression from the 2:1 and 1:1 releases (57.6% and 54.6% mean hatching rate respectively, nonsignificant), compared to the 0:1 control (72.4% mean hatching rate) (**Figure 2B,C**). Release ratios of 10:1 or higher are common (63) for other sterile transgenic mosquito control campaigns, demonstrating that *A. gambiae* pgSIT males achieve sufficient suppression to be considered candidates for SIT releases, but larger trials are needed.

**Figure 2:**
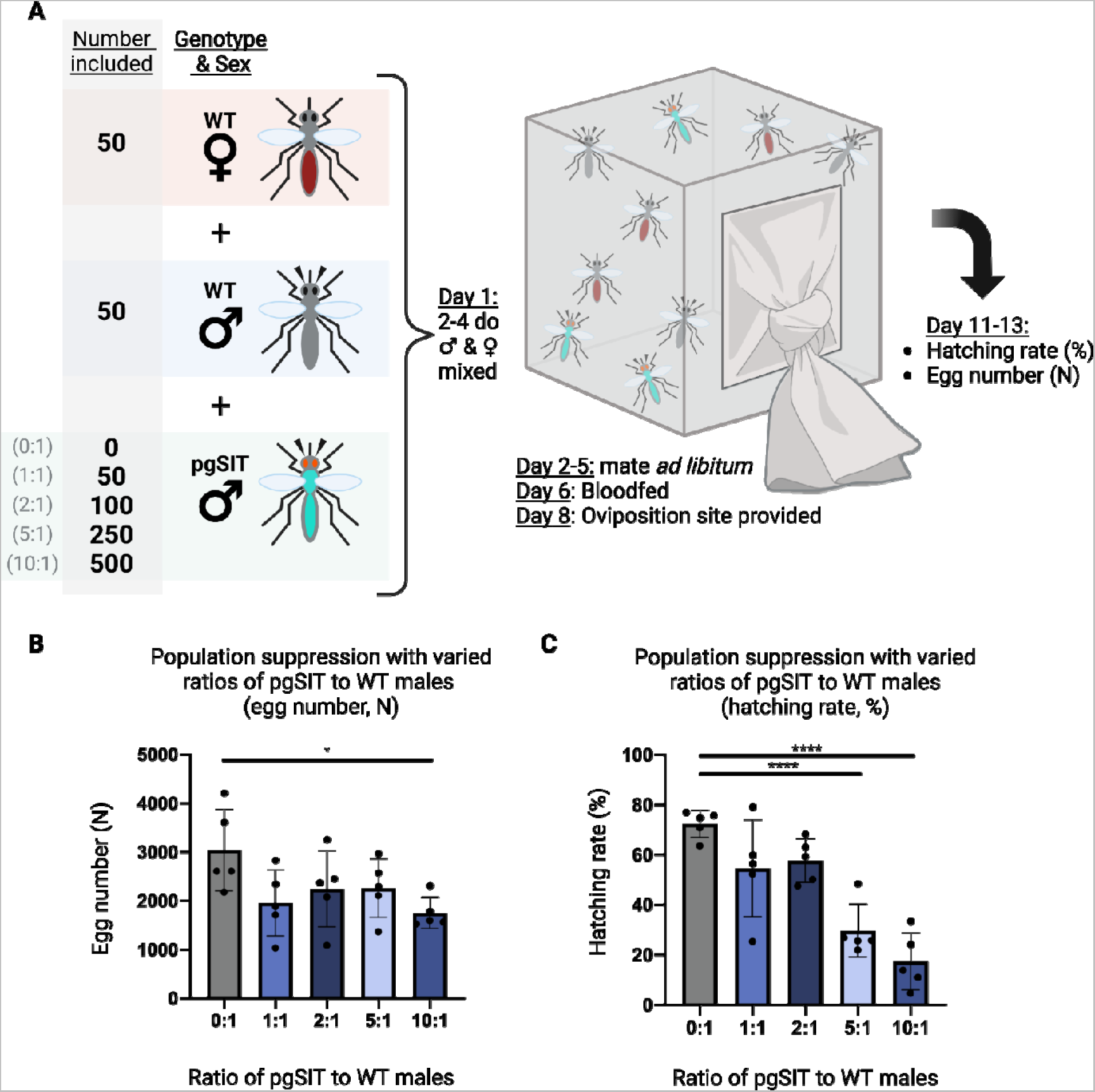
Population suppression following release of pgSIT males at different ratios to wild type. **A**) Test suppression cages were established with 50 wild type males, 50 wild type virgin females, and either 0, 50,100, 250, or 500 pgSIT^D15^ males (for the 0:1, 1:1, 2:1, 5:1, and 10:1 pgSIT:wild type male ratios respectively). After mating and blood feeding, the hatching rate was calculated for each cage. **B)** The egg counts from population suppression assay cages. Groups 0:1 and 10:1 are significantly different (p < 0.05, Dunnett’s multiple comparisons). Mean and SD shown. **C)** Population suppression as measured by the hatching rate (%) from cage suppressed by different ratios of pgSIT males to wild type males. Hatching rate is reported as the percent of eggs which hatched (n% = n 1 day old larvae / n eggs laid). The 0:1 control group differs significantly with both the 5:1 (p < 0.001) and 10:1 (p < 0.001) groups (One Way ANOVA, Dunnett’s multiple comparisons test). Mean and SD shown.

The broods from population suppression assays were also monitored for the presence of fluorescent transgenic F2 offspring which would indicate a fertile pgSIT^D15^ father. Among the 20 cages tested containing pgSIT males (**Figure 2B,C, Table S6**), only a single brood yielded transgenic larvae (n = 43 transgenic larvae total, from the 2:1 suppression group), suggesting the presence of a single fertile male which escaped the sterilization phenotype, providing evidence of the only fertile pgSIT male observed throughout the course of this work.

### Modeling pgSIT as a suppression technology

We next modeled hypothetical releases of pgSIT *A. gambiae* eggs to explore their potential to eliminate a local *A. gambiae* population resembling that of the Upper River region of The Gambia using the MGDrivE 2 framework (64) with parameters listed in **Table S7**. The modeling framework was calibrated to malaria prevalence data from a randomized controlled trial conducted in the Upper River region (65), and informed by local entomological data (66) and rainfall data sourced from Climate Hazards Group InfraRed Precipitation with Station data (CHIRPS, https://www.chc.ucsb.edu/data/chirps). Parameters describing the pgSIT system were based on results from this paper suggesting the pgSIT system in *A. gambiae* induces near complete male sterility (>99.5%) and female inviability (>99.9%), with inviability being modeled at pupation (allowing inviable individuals to contribute to density-dependent larval mortality without mating as adults). To be conservative, we assumed a 25% reduction in both pgSIT male mating competitiveness and lifespan compared to wild-type males, as fitness costs sometimes emerge in the field (63), despite no reductions in lifespan being observed in this work.

With the parameterized modeling framework in place, we simulated 0-20 consecutive weekly releases of pgSIT eggs at a ratio of 0-160 pgSIT eggs (female and male) per wild *A. gambiae* adult (female and male) (**Figure 3**). Previous pgSIT modeling studies (52, 53) suggested that *Aedes aegypti* populations could potentially be eliminated by 10-24 consecutive weekly releases of 40-400 pgSIT eggs per wild adult; however, we focused on release schemes having smaller weekly release sizes as a more cost-effective option. The mosquito population in the Upper River region is highly seasonal, as reflected in the first two years of the time-series dynamics (pre-release), so we simulated pgSIT eggs released from the beginning of the rainy season (June 1st), just as the *A. gambiae* population begins to grow - a timing determined optimal for several genetic control systems (63, 67). Simulation output predicts local *A. gambiae* elimination for achievable release schemes - ≥14 weekly releases of 40 pgSIT eggs per adult mosquito, ≥11 weekly releases of 80 pgSIT eggs per adult mosquito, and ≥9 weekly releases of 128 pgSIT eggs per adult mosquito. In many cases where elimination is not achieved, significant population suppression still occurs and is maintained for >6 months, which would be expected to have a significant epidemiological impact.

**Figure 3.**
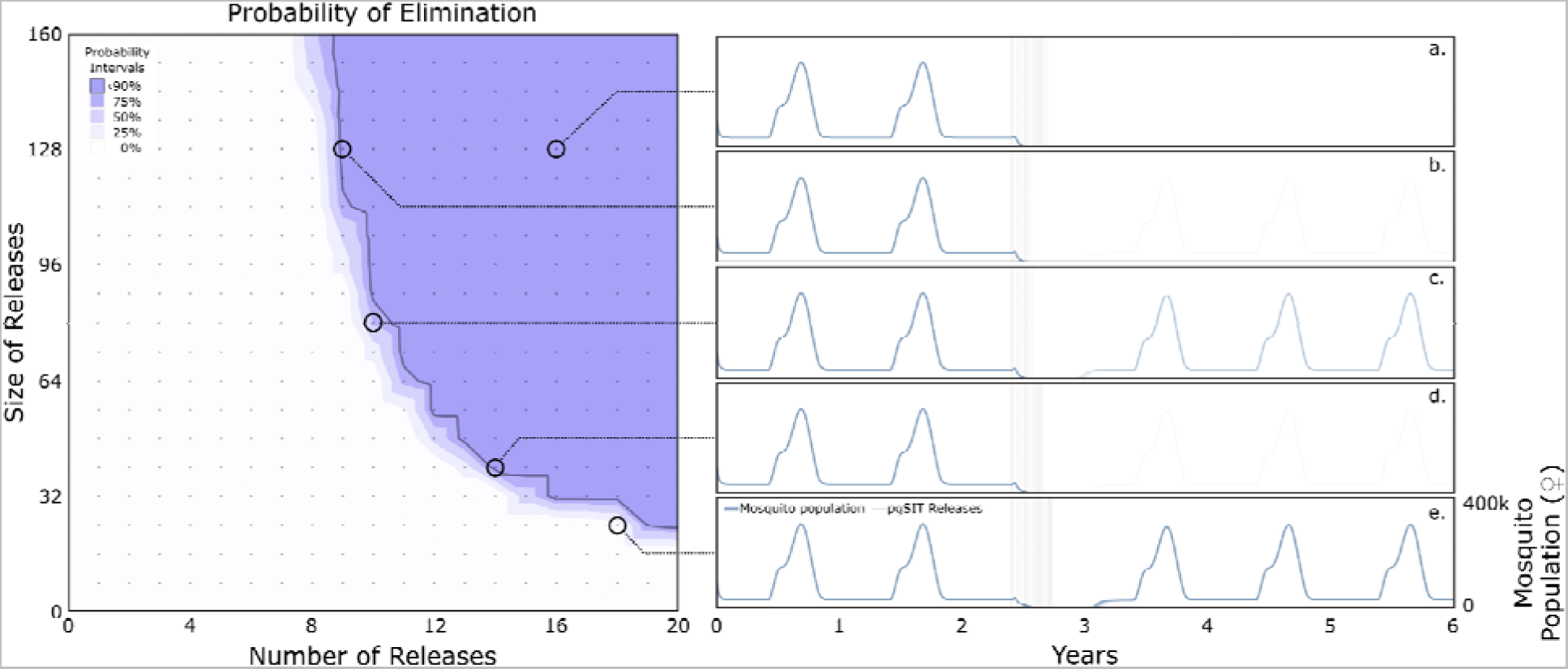
Modeling population suppression and elimination by release of pgSIT *A. gambiae*. Weekly egg releases were simulated in a randomly-mixing *A. gambiae* population resembling the Upper River region of The Gambia using the MGDrivE 2 simulation framework ^66^ with parameters described in **Table S8**. Probability of *A. gambiae* population elimination is depicted for a range of release schemes described by the number of consecutive weekly releases and number of pgSIT eggs released per wild-type adult. The contour plot (left) depicts regions of parameter space for which the local mosquito population is eliminated with probabilities ≥0, 25, 50, 75 or 90% (as measured by the proportion of 100 simulations that lead to elimination). 90% elimination probability is depicted by a solid line. Time-series mosquito population dynamics (right) are depicted for a selection of scenarios from the contour plot. Releases of pgSIT eggs (female and male) are modeled beginning June 1st (beginning of the Upper River rainy season) in the 3rd year of the simulation. Scenario (a) depicts a large release very likely to achieve elimination (16 weekly releases at a 128:1 ratio of pgSIT eggs to wild adults); scenarios (b) and (d) depict release schemes ~90% likely to achieve elimination (9 weekly releases at a 128:1 ratio, and 14 weekly releases at a 40:1 ratio, respectively); scenario (c) represents a release scheme with a ~75% elimination probability (10 weekly releases at a 80:1 ratio), with the population rebounding in ~25% of simulations (faint lines in years 4-6); and scenario (e) represents a release scheme that achieves transient suppression but not elimination.

## Discussion

In this work, we develop the genetic SIT technology, pgSIT, in the malaria vector *A. gambiae*, meeting the demand for a confinable and proven mosquito suppression technology in this species. Overall, we demonstrate that our pgSIT system exhibits remarkable male sterilization (100% in assays, >99.5% for the population as a whole), and female elimination (100% in assays, >99.9% for the population as a whole), and strong population suppression effects in cage trials, yielding a system amenable to high-throughput safe SIT releases of pre-sterilized and pre-sex-sorted eggs. In detail, we generated a novel gRNA-expressing transgene, gZBD, targeting *zpg, dsxF,* and β*2-tubulin* for CRISPR cleavage. When crossed to a Cas9 transgenic line, we observed in the hybrid progeny significant but incomplete female androgenization due to *dsxF* targeting. We also observed complete or nearly complete sterility of the (+/gZBD^A18^; +/Cas9) or (+/gZBD^D15^; +/Cas9) respectively, due to *zpg* and β*2-tubulin* targeting. To improve female-elimination we crossed the gZBD line to the Ifegenia Genetic Sexing System line, gFLE and established double homozygous gRNA-expressing lines (gFLE;gZBD^A18^) and (gFLE;gZBD^D15^). When crossed to Cas9 we confirmed the presence of mutations within each gene in the hybrid progeny, however crosses generated with the Cas9 transgene derived maternally were lethal. While further elucidation of this phenotype was beyond the scope of this work, we postulate this is due to an overabundance of on- and off-target cleavage because maternal Cas9 is expected to yield stronger embryonic mutagenesis due to maternal deposition by the Vasa promoter (68). Remarkably, pgSIT individuals of both lines exhibit strong female-elimination, 99.8% and 100% respectively for pgSIT^A18^ and pgSIT^D15^, in addition to high levels of sterility, >99.5% each. From these two lines, we selected pgSIT^D15^ for more in depth characterization. In line with the sterility phenotype, pgSIT^D15^ males lacked testes but maintained otherwise normal Lower Reproductive Tracts (**Figure S4A**). Interestingly, (gFLE/Cas9) controls also displayed aberrant and occasionally absent testes, a phenotype not noted in prior work due to population fertility as a whole (69). Though further elucidation is beyond the scope of this work, we postulate that it’s mimicking the function of *fle*’s closest well characterized homolog, *Transformer2* (*tra2*), whose misregulation during fly spermatogenesis causes infertility and defective sperm (70). This finding potentially explains why the pgSIT^D15^ individuals in (**Figure 1D**) had higher rates of sterility than the (gZBD^D15^/Cas9) males in **Figure 2C**, if this phenotype has an additional sterilization effect. In line with having an otherwise normal reproductive tract, we confirmed that pgSIT males were able to transfer a mating plug, a key requirement for induction of refractoriness in females (**Figure S4B**)(60, 61).

We further confirmed that pgSIT^D15^ males are long-lived by Survival Curve Assay, and validated their genomic integration locus by Nanoopore. Interestingly, Nanopore also revealed a deletion of the gRNA^dsxF.1^ expression cassette while confirming the presence of the other gRNAs. This was confirmed by Sanger sequencing, potentially explaining some findings throughout this work. While clear intersex phenotypes were observed in early assays (**Figure S1**), they were partially penetrant (**Table S1**). In later assays, intersex phenotypes were completely lost (siblings of **Figure S2**) likely after gRNA^dsxF.1^’s deletion, concurrently revealing that mutagenesis by the remaining *dsxF*-targeting gRNA, gRNA^dsxF.2^, does not cause intersex phenotypes. We postulate this mutant expanded within the population following population bottlenecks caused by the COVID-19 pandemic, concurrently raising concerns about the long-term stability of repetitive multi-gRNA transgenes including some gene drives (8, 71). Though interesting, female-killing by gFLE makes targeting *dsx* redundant, rendering this deletion inconsequential to pgSIT function. In all, although the genetics is not as originally designed, the pgSIT system still produces all of the desired necessary phenotypes to be candidates for further trials. In competition cage trial assays, we demonstrate that pgSIT^D15^ males are capable of causing significant population suppression when competing against wild type males at 10:1 and 5:1 ratios (both p < 0.0001, Dunnett’s multiple comparison) yielding a 4.2x and 2.4x fold reduction in average hatching rate respectively, a strong suppression phenotype similar to those observed in pgSIT in other organisms (52, 53). Finally, modeling suggests this system is capable of eliminating local *A. gambiae* populations, and hence interrupting malaria transmission, for achievable release schemes of ≥14 releases of 40 pgSIT eggs per adult mosquito or ≥11 weekly releases of 80 pgSIT eggs per adult mosquito. In total, this work demonstrates that this pgSIT system exhibits all of the necessary properties for consideration as a releasable line for SIT-like vector control of *A*. *gambiae*.

The pgSIT system outlined here may also enable suppression of the adjacent species within the *A. gambiae* complex: *A. arabiensis*, *A. quadriannulatus*, *A. melas*, and *A. merus.* Not only are the target sites for transgenic *gRNAs,* gRNA^zpg.1^, gRNA^B2.2^, gRNA^dsxF.1^, gRNA^dsxF.2^, gRNA^fle.7^, and gRNA^fle.10^ conserved making introgression into these species possible (72, 73), but an overabundance of released *A. gambiae* pgSIT males may breach natural mating barriers to directly suppress these species as well (74–77). With gene drives being proposed to spread beyond target species assuming the drive target site is conserved and not-mutated (78), the possibility of this occurring with other vector control strategies such as pgSIT should be explored as well.

Compared to other vector control methods, pgSIT is more scalable and can be released during all life stages. For GM vector control campaigns except RIDL and sex-biasing gene drives (6, 79), the rate-limiting step for releases is sorting males from the undesirable, disease-transmitting, females. If not performed manually, which is limited to 500 pupae per hour (80), this is achieved by optical sorting or by fluorescence-based sex-sorters (46, 48, 81–83). The former utilizes an AI-trained camera to distinguish between sexes as adults (82, 83), while the latter relies on transgenic sex-specific fluorescence to enable sorting of nascent larvae by COPAS (49). For other vector control measures, sorting occurs on the individuals directly to be released, yielding a fairly low 2:1 sort:release ratio (**Text S1**) (29, 39, 40). In pgSIT however, because sorting occurs the generation prior to release (F0 generation), and the released generation is automatically sex-sorted (F1 generation), pgSIT, and the sister technology Ifegenia have a sort:release ratio closer to 1:50 (**Text S1**). These features not only make pgSIT higher-throughput by orders of magnitude, but also enable delivery of eggs via drone (84, 85). While manual F0 sorting was performed in this study, a technique effective for small scale field trials (86), optical F0 sorting could enable immediate commencement of larger-scale trials and possibly even release campaigns. Future iterations of pgSIT, termed pgSIT 2.0, could consolidate the existing transgenes and include fluorescent-sex sorters to further improve scalability (87). Maximizing throughput in pgSIT2.0 could yield 40,000 COPAS-sorted F0 larvae per hour from a single machine (49), yielding 2 million F1 sterilized males in the next generation, potentially facilitating production and releases on a continental scale (73).

While pgSIT does not aim to release transgenes into the population, our observation of rare fertile escapee males indicates that release of some CRISPR transgenes into the population will likely occur. It has been shown that population eradication by pgSIT does not require complete (100%) sterility penetrance, as appreciable levels of suppression can be achieved by incompletely penetrant systems (55). The released transgenes would separately express Cas9 and gRNAs, but they are incapable of gene drive given their dislinkage and genomic position, and are expected to be lost from the population given their inherent fitness defects. Importantly, these alleles may include rare resistance alleles, however because fresh pgSIT individuals would be released iteratively, wild females carrying these alleles would be sterilized and prevented from transmitting it further to their offspring.

As a confineable and scalable vector control technology, pgSIT is a valuable addition to the *A. gambiae* vector control toolkit. Previous culicinae-based technologies have either been not yet developed in this species, are not confineable, not scalable, or release potentially undesirable transgenes into the population; RIDL, Gene Drives, X-shredders and Ifegenia respectively. One powerful confinable, scalable, and field-proven GM technology is RIDL, however its chemically-repressed daughter-incapacitating effects have not yet been developed in *A. gambiae* (63). Among those developed in *A. gambiae*, the most advanced and scalable technology are gene drives (6, 88), but their uncontrollability has raised political, ethical, ecological and socio-economic concerns, hindering their release (14). The long term durability of drives is unclear as many generate their own resistance mutations due to the constitutive co-expression of CRISPR components, ultimately hindering their spread. Following release of population suppression drives, selection pressures for resistance mutations which evade extinction will be very high making the system prone to breakage, as they rely on the integrity of a single gRNA target site. That said, pgSIT is quite robust as the two CRISPR strains are maintained separately, preventing resistance allele generation, and only producing the released ‘dead end’ males following a controlled cross in the production facility. On the other hand, X-shredders are a confinable alternative in *A. gambiae*, but lack scalability. By targeting the X-chromosome for shredding, this technology achieves male-biasing or offspring-killing, but lacks an inducibility feature making control of the phenotypes problematic. To scale for recent field trials (86) manual sorting for both the transgene and for the sex yielded a sort to release ratio greater than 2:1 making it unsuited to larger scale trials at this time. Finally, a non-driving scalable suppression technology, termed Ifegenia, was recently published in *A. gambiae* (51) which uses a binary CRISPR crossing system similar to pgSIT to induce multigenerational female killing. While Ifegena has an identical sort:release ratio as pgSIT (1:50, **Test S1**), it results in the persistence - though not drive - of GM CRISPR transgenes in the population which may be undesirable for some applications. Taken together, pgSIT fills a unique niche between these technologies; it is more high-throughput than X-shredders, more confinable and controllable than gene drives, and because it is a “genetic dead end” it does not release transgenes into the population like Ifegenia, making it a powerful and highly scalable new tool in the *A. gambiae* vector control toolkit.

In all pgSIT presents a novel advancement for vector control of *A. gambiae.* It exhibits all criteria for a target product profile for use in SIT releases; it produces highly sterile males and in mass, it displays advantageous fitness parameters, and it is more confinable and more scalable than alternative GM technologies. In all, pgSIT presents a powerful new tool in the toolkit for control of this deadly malaria vector, potentially revolutionizing control of this deadly pest.

## Methods

### Mosquito rearing and maintenance

*A. gambiae* used in this work was derived from the stock G3 strain. Mosquitoes were reared in an ACL-2 insectary under 12hr light/ dark cycles at 27°C with a water source provided for drinking and ambient humidity. Adult mosquitoes were placed in Bugdorm, 24.5LJ×LJ24.5LJ×LJ24.5LJcm cages. Adults were fed 0.3 M aqueous sucrose *ad libitum*. Males and females were allowed to mate for 4-7 days prior to being provided with a blood meal on anesthetized mice for about 15 minutes. Egg dishes, composed of urinalysis cups filled with water and lined with a filter paper cone, were provided to cages 48 h after a blood meal. Eggs were allowed to melanize and hatch unperturbed for 3 days in the egg dish before being floated into trays filled with DI water. Larvae were reared and fed, and pupae were screened and sexed, in accordance with established protocols (89).

### Cloning and plasmids

Cloning and molecular biology work was undertaken using established cloning protocols. Plasmid 1114H (gZBD) is available at Addgene (200640). All other transgenes used in this work were previously published (51, 58)

### gRNA design

The target gene reference sequences for *zpg* (AGAP006241), *B2-tubulin* (AGAP008622), *dsxF* (AGAP004050) were retrieved from VectorBase (90), and sequences were confirmed by PCR. gRNA’s were designed using software available at http://crispor.tefor.net. Two gRNAs targeting *dsxF* were selected, one from the literature, gRNA^dsxF.1^ (5’ GTTTAACACAGGTCAAGCGG 3’) (6), and the second designed *de novo* to target the extreme 3’ end of the *dsxF* exon 5 coding sequence, gRNA^dsxF.2^, (5’ TTATCATCCACTCTGACGGG 3’). Two gRNAs targeting the first exon of *B2-tubulin* were designed, gRNA^B2.1^, (5’ GCTCGATATCGTGCGCAAGG 3’), and gRNA^B2.2^, (5’ CCAAATAGGCGCTAAGTTCT 3’). A single gRNA targeting the first exon of *zpg* previously shown to cause robust germline mutagenesis (41, 58, 91) was used, *gRNA^zpg.1^* (5’ GATCCGATCACGCAGTCGAT 3’). The gRNAs gRNA^fle.7^ (5’ CGACGGCTCGTTCATCGCTG 3’) and gRNA^fle.10^ (5’ ATCGAGCGCGTCGCCTGGTA 3’) targeting *fle* were previously described (51, 92).

### Embryonic microinjections

Injections were carried out as described previously (93–95). In brief, the gZBD plasmid injection mix was prepared by maxiprep and was diluted to prepare 100µl of a 350 ng/µl solution in diH2O. 45m - 2h old embryos were harvested from a stock cage of the G3 line and aligned on a glass slide, posterior end up, along the edge of a dampened Millipore mixed cellulose esters membrane (CAT No. HAWP04700F1) covered with a cut-to-size Whatman filter paper (CAT No. 1001-150), as diagrammed in (93). The posterior end of embryos were injected with a quartz needle filled with injection mix and controlled by an Eppendorf FemtoJet4x injection system (CAT No. 5253000025). Injected embryos remained on the slide, and the slides were placed in a water dish with the end of the Whatman filter paper submerged to permit capillary action to prevent the eggs from drying. Neonate F0 larvae were removed beginning 48h post injection and were reared separately.

### Fluorescent Sorting, Sexing and Imaging

All imaging of *A. gambiae* was carried out under a Leica M165FC fluorescent stereomicroscope outfitted with a Leica DMC2900 camera. Fluorescence was visualized using the CFP/YFP/mCherry triple filter, and pupal sex was determined by examining the pupal genital terminalia as diagrammed in (96).

### gZBD family establishment

The nomenclature for the F0, F1, and F2 generation demarcations within this section of the methods follows the more traditional use of these generational markers within the field. It differs from the use of F0 and F1 in the main text which is in reference to stock parental (F0) and hybrid (F1) generations for study of pgSIT. They are in reference to different experiments and genotypes, and are not to be confused.

To establish gZBD transgenics, embryonic microinjection of the gZBD transgene was carried out into F0 individuals essentially as described above. F0’s were reared to adulthood, outcrossed to wild type G3 stock line of the opposite sex, and blood fed. The resulting F1 offspring yielded multiple F1 ‘founder’ transgenic larvae which were identified by fluorescence. Female F1 individuals were isolated individually and used to found the gZBD^A^ and gZBD^D^ families. The gZBD^A^ and gZBD^D^ families both exhibited fluorescence patterns indicative of multiple insertion sites. Therefore, to generate sub-families with single insertion sites, gZBD^A^ and gZBD^D^ female transgenics were outcrossed to G3 wild type males in bulk (over 100 individuals of each sex) for 5 generations, selecting for female transgenics each generation. Single females from the ‘diluted’ gZBD^A^ and gZBD^D^ lines were isolated and allowed to lay separately, and a single brood that had uniform fluorescent patterns suggesting a single insertion site from each family was used to found the gZBD^A18^ and gZBD^D15^ subfamilies.

### Identifying and validating genomic insertion site of gZBD transgenes

To identify the genomic insertion site of gZBD, genomic samples were taken from crushed gZBD^A18^ and gZBD^D15^ adults, and inverse PCR was performed essentially as described in (97). In short, 1-3 μg of genomic DNA were treated with TaqI restriction enzyme for 4h, then circularized with ligase in a dilute 100μl reaction. The sample was re-concentrated by Sodium Acetate precipitation followed by resuspension in 10 μl water, of which 1μl was used to template the inverse PCR. PCR was carried out with the primers 1114H.S3 and 1114H.S4, (5’ CTGTGCATTTAGGACATCTCAGTC 3’) and (5’ GACGGATTCGCGCTATTTAGAAAG 3’) respectively, the latter of which amplifies outwards beyond the piggyBac terminal repeat of the gZBD plasmid and into adjacent genomic sequences. PCR amplicons were gel extracted, cloned into pJET (Thermo Scientific, Cat. No. / ID: K1231), and individually sequenced. Reads were aligned against the piggyBac terminal repeat of the gZBD transgene with all sequencing beyond the repeat terminus corresponding to the locus of integration. Through this method, gZBD^A18^ and gZBD^D15^ were found to be integrated in the (chr3L:34188038) and (chr3L:828896) loci respectively. Primers for standard PCR were designed to confirm the genomic integration, and used to homozygous the transgenic lines used throughout this work. To identify the transgenic gZBD^A18^ allele, the primers 1114H.S4 and 1114H.ipS1; (5’ GACGGATTCGCGCTATTTAGAAAG 3’) and (5’ CATTGAACGGTCTATGCTGTCATGTAC 3’) respectively, were used for PCR amplification. To identify the presence of the wild type (unintegrated) gZBD^A18^ allele, the primers 1114H.ipS1 and 1114H.S31, (5’ CATTGAACGGTCTATGCTGTCATGTAC 3’) and (5’ CGTTCTTGCGAAAAGGTGAAAAGTG 3’) respectively, were used. To identify the transgenic gZBD^D15^ allele, primers 1114H.S17 and 1114H.S29, (5’ GACTGAGATGTCCTAAATGCAC 3’) and (5’ CTCGTGACCCTCGTTATAG 3’) respectively, were used, while the primers 1114H.S30 and 1114H.S29, (5’ CATGTTGTTCTTTTGGAAAGC 3’) and (5’ CTCGTGACCCTCGTTATAG 3’) respectively, were used to identify the presence of a wild type (unintegrated) gZBD^D15^ allele.

### **Δ***dsx* knockout phenotype characterization of gZBD families

Following embryonic microinjections of gZBD into F0 embryos, the F1 generation yielded transgenic ‘founder’ larvae. While female F1 founders were used to establish clonal iso-female lines for study, the male F1 founders - with mixed uncharacterized and unknown insertion sites - were crossed to Cas9 females in bulk. The resulting F2 transheterozygous hybrids (+/gZBD; +/Cas9) were imaged for genital androgenization (**Figure S1**).

### Male sterility characterization of +/gZBD and +/gFLE;gZBD families

For crosses assaying male sterility, we established cages of 50 transgenic hybrid sterile males - (+/gZBD; +/Cas9) or pgSIT (+/gFLE;gZBD; +/Cas9) - to 50 virgin wild type females on day 1, and allowed them to mate *ad libitum*. On day 6 females were fed a mouse blood meal, and an oviposition site (egg dish) was provided on day 8. Larvae were counted on days 11, 12, and 13 and checked for the presence of fluorescence. If F2 larvae were fluorescent at transgene ratios expected of progeny from a hybrid transgenic father, they were counted and presumed to belong to an escapee fertile male. These F2 offspring for (+/gZBD; +/Cas9) sterility experiments were collected and sequenced for mutations at the gRNA target sites following sequencing protocols listed above. If F2 larvae were completely non-fluorescent, then a contamination was presumed to have occurred via inclusion of a non-transgenic male, and the replicate discarded (one replicate). Eggs and eggshells were counted on days 13 and 14. Hatching rate was calculated as the number of larvae over the number of eggs (**Figure 1C**, **Figure S2B**), the number of eggs is also reported (**Figure 1D** and **Figure S2C**).

### Establishing homozygous pgSIT^A18^ and pgSIT^D15^ lines

To establish the doubly homozygous gFLE;gZBD^A18^ and gFLE;gZBD^D15^ lines, we began by crossing gZBD^A18^ and gZBD^D15^ separately to gFLE. For five generations brightly fluorescent individuals with an ‘aqua’ fluorescence color, indicative of dual *EGFP* and *m2Turquoise* fluorescence, were sorted for as pupae and allowed to mate *ad libitum*. Then on the fifth generation, individuals were fluorescently sorted and allowed to mate *ad libitum* as described above, but following blood feeding females were isolated into single oviposition cups to lay egg clutches in isolation. From each resulting brood, a small pool of individuals were taken as L1 larvae to check for gFLE and gZBD homozygosity via PCR. Primers 1154A.S32 and 1154A.S3, (5’ CTTTCTAACGGTACGCAGCAG 3’) and (5’ AACAGCCACAACGTCTATATCATG 3’) respectively, were used to identify the presence of the transgene in the gFLE transgenic locus, while the primers 1154A.S32 and 1154A.S34, (5’ CTTTCTAACGGTACGCAGCAG 3’) and (5’ GCTCCAGTTCATGTCGATAGAC 3’) respectively, were used to identify the presence of a wild type gFLE locus. Primers for analysis of gZBD^A18^ and gZBD^D15^ loci are listed above (see **Identifying and validating genomic insertion site of gZBD transgenes** in **Methods**).

### Crosses to generate F1 gRNA/Cas9 hybrids

For all crosses, pupae were fluorescently sorted and sexed, and allowed to emerge as adults in separate cages to ensure female virginity before crossing. Unless otherwise indicated, crosses of 50 males x 50 females were set up on Day 1 with 2-4 day old adults, allowed to mate *ad libitum*, then blood fed on day 6. The crosses to generate the (+/gZBD; +/Cas9) genotype in **Figure S1** were generated with maternal Cas9 paternal gRNA F0 directionality, while the crosses to generate the same genotype of males in **Figure S2** used the reciprocal cross. The crosses to generate the F1 pgSIT (+/gFLE;gZBD; +/Cas9) genotype were performed with F0 Cas9 females and gRNA males for mutation analysis in **Figure S3**, but following identification of the lethal phenotype of this cross directionality, all subsequent crosses to generate this genotype used the Cas9 male and gRNA female directionality (**Figure 1C,D**, **Figure 2B,C**).

### Quantifying female elimination of F1 (+/gFLE;gZBD; +/Cas9)

F1 gRNA/Cas9 hybrids generated with maternal gRNA and paternal Cas9 were sex-sorted daily as pupae. Counts of males and females were recorded starting the first day pupation is observed until the day all larvae had become pupae (**Figure 1B** and **Table S4**), typically 4-6 days. Male and female pupae were placed in separate cages to emerge as adults, and the survival of female pupae was closely monitored. Females who emerged as adults and were able to fly were crossed to 50 wild type adult males and allowed to mate *ad libitum*, observed during blood feeding for their ability to take blood meal, and given an egg dish 48 hrs post blood feed.

### Testes dissections

4-day-old adult virgin males were immobilized on ice for <1h, and the lower reproductive tract was dissected into PBS by pulling slowly from the claspers. Images were taken with a Leica M165FC fluorescent stereomicroscope outfitted with a Leica DMC2900 camera under 6.5x magnification. Lighting orientation, brightness, exposure time, and white balance were not controlled for, so no conclusions about tissue color, brightness, or tone should be made from these images (**Figure S4A**).

### Mating plug transfer assay

To determine whether or not pgSIT males could transfer a mating plug, we crossed 100 pgSIT^D15^ or wild type 5-7 day old virgin males to 100 wild type virgin 5-7 day old females at dusk when males were swarming. We allowed them to mate *ad libitum* for 45 minutes, during which we verified the presence of copulating pairs at the bottom of the cage. After 45 minutes, all females were removed onto ice. The terminal abdominal segments of females were imaged, ventral side up, with a Leica M165FC fluorescent stereomicroscope and a CFP/YFP/mCherry triple filter (**Figure S4B**). The presence of the mating plug could be seen through the female cuticle by autofluorescence within the female atrium - a previously established assay for verifying mating plug transfer (60, 98). Lighting orientation, brightness, exposure time, and white balance were not controlled for, so conclusions about plug brightness should not be made from these images.

### Male adult survival assay

17 male pupae were put into each small Bugdorm cage on day 0. This number of males was selected to minimize crowding and competition between males. On day 1, the number of dead pupae or drowned adults were counted and removed, but were not included in survival curve counts. From day 2 onward the number of adult dead males were counted, removed, and recorded each day. All cages within a replicate were summed for the final survival curve analysis, yielding a total of 110 wild type males and 81 pgSIT^D15^ males analyzed. At the end of the assay, for cages that had no more living mosquitoes but had individuals that were unaccounted for (4 cages out of 12 cages total), the unaccounted individuals were censored on the final day of the survival curve analysis and marked as censorship notches in **Figure S4C**. Raw survival counts broken down by cage can be found in **Table S6**.

### Insertion site mapping

Insertion sites for gFLE and Cas9 transgenes were previously determined to be located at 2R(NT_078266.2): 23,279,556-23,279,559 and 2L(NT_078265.2):10,326,500-10,326,503, respectively (51). To determine insertion site for the gZBD transgene, we performed Oxford Nanopore sequencing of genomic DNA from adult transheterozygous pgSIT^D15^ males harboring the +/gZBD transgene in addition to the +/Cas9 and +/gFLE transgenes. DNA was extracted in pools of 6-8 adult mosquitoes mosquitoes using Blood & Cell Culture DNA Midi Kit (Qiagen, Cat# 13343) following the manufacturer’s protocol. The sequencing library was prepared using the Oxford Nanopore SQK-LSK110 genomic library kit and sequenced on a single MinION flowcell (R9.4.1) for 72 hrs. Basecalling was performed with ONT Guppy basecalling software version 6.4.6 using dna_r9.4.1_450bps_sup model generating 3.92 million reads above the quality threshold of Q≧10 with N50 of 6608 bp and total yield of 14.43 Gb.

To identify transgene insertion sites, nanopore reads were aligned to the gZBD plasmid sequence (Plasmid #1114H, Addgene #200640) using minimap2 (99). Reads mapped to the plasmids were extracted and mapped to *A. gambiae* genome (GCF_000005575.2_AgamP3). Exact insertion sites were determined by examining read alignments in Interactive Genomics Viewer (IGV). The gZBD transgene is integrated between positions 4,828,892 and 4,828,896 on chromosome 3L (NT_078267.5). The site is located in the intergenic region between AGAP010485 and AGAP010486. The previously determined integration sites for gFLE and Cas9 transgenes were confirmed with the new nanopore data. The nanopore sequencing data have been deposited to the NCBI sequence read archive (PRJNA978105)

### Sequencing of gRNA expression cassettes

gDNA from gZBD^D15^ and pgSIT^D15^ were extracted (Qiagen, DNeasy Blood & Tissue Kits, Cat. No. / ID: 69504) from pools of 3 adults, PCR amplified (Q5 HotStart DNA polymerase (NEB, Cat. No./ID: M0493L)), and Sanger sequenced for the 7 gRNA expression cassettes. The gRNA^dsxF.1^ expression cassette was amplified and sequenced with the 1114E.S14 (5’ CCCGTCAGAGTGGATGATAAC 3’) and 1114A.S16 (5’ GCTTACGTTTACTGCTATCTGCACTTC 3’) primers. The gRNA^dsxF.2^ cassette was amplified and sequenced with the 1114H.S7 (5’ CGGTTTTGTTTGCAGCGAGTTGTG 3’) and aa151 (5’GGTAATCGATTTTTTCAGTGCAG 3’) primers. The gRNA^zpg.1^, gRNA^B2.1^, gRNA^B2.2^, gRNA^fle.7^, and gRNA^fle.10^ expression cassettes were amplified and sequenced all together with the 1114H.S1 (5’ CTCAAAATTTCTTCTATAAAGTAACAAAAC 3’) and 1114G.C2 (5’ CGAGGTTCTCCTTATGCTCTGTG 3’) primers. PCR amplicons were run on 1% agarose gel at 120V for 20 minutes, then gel extracted with the Zymoclean Gel DNA Recovery Kit (Zymo Research, Cat. No./ID: D4007).

### Target site mutation analysis

Mutations under the gRNA target sites were identified in F1 (+/gFLE;gZBD^D15^; +/Cas9) hybrid offspring resulting from a cross between 50 (gFLE;gZBD^D15^) males and 50 Cas9 females. The male-female directionality of this F0 cross was chosen because Cas9 females provide maternal deposits of Cas9 protein into the embryo, producing F1 hybrid offspring with a high mosaic mutation load and allowing for sequencing of many mutant alleles. F1 hybrid offspring were collected in bulk as late-stage embryos and were DNA extracted (Qiagen, DNeasy Blood & Tissue Kits, Cat. No. / ID: 69504) and PCR amplified (Q5 HotStart DNA polymerase (NEB, Cat. No./ID: M0493L)). The *zpg* locus was amplified with the 114H.S34 and 1114H.S37; 1114H.S34 (5’ GTAGAAAGAGCAAGGAAAGAAACG 3’) and 1114H.S37 (5’ GTTCCGAATTTCCAAGTGCTTC 3’) primers respectively. The β*2-tubulin* locus was amplified with the 1114H.S38 (5’ GCTAAATATCAGACGGCTTTC 3’) and 1114H.S39 (5’ GCGAATTTTCGAAATCAGCAG 3’) primers. The *dsxF* locus was amplified with the 1114E.S33 (5’ CTTGCCATCCTATGGAACTGC 3’) and 1114E.S32 (5’ GGTGAAAATATTGTTGATGCGC 3’) primers. The *fle* locus was amplified with the aa174 (5’ CGACTCACTATAGGGAGAGCGGC 3’) and aa175 (5’ AAGAACATCGATTTTCCATGGCAG 3’) primers (51). PCR amplicons were run on 1% agarose gel at 120V for 20 minutes, then gel extracted with the Zymoclean Gel DNA Recovery Kit (Zymo Research, Cat. No./ID: D4007). Purified amplicons were then cloned into the pJET vector (Thermo Scientific, Cat. No. / ID: K1231), transformed into chemically competent *E. coli* (Promega, JM109), and plated on LB-Ampicillin plates. Plates were sent for Sanger Colony sequencing with universal primers PJET1-2F (5’ CGACTCACTATAGGGAGAGCGGC 3’) and/or PJET 1-2R (5’ AAGAACATCGATTTTCCATGGCAG 3’), with each colony representing a single PCR amplicon from an individual mutant allele.

Because their mutation frequency was qualitatively weaker, to enrich for mutant alleles under gRNA^B2.1^, gRNA^B2.2^, gRNA^dsxF.1^, and gRNA^fle.7^, their genomic target sites were PCR-amplified, and these PCR amplicons were digested with a restriction enzyme whose recognition site overlaps the expected gRNA cut sites, such that an undigestible PCR product indicates a likely CRISPR mutation (**Figure S3**) (100). The β*2-tubulin* locus was amplified with the 1114A.S43 (5’ GAGAGCAACACTCGTGCG 3’) and 1114A.S44 (5’CAGGGTGGCATTGTACG 3’) primers and the amplicon was digested with Fspl (NEB cat#R0135S) or Ddel (NEB cat#R0175S) to identify mutations by gRNA^B2.1^ and gRNA^B2.2^ respectively. To identify mutations by gRNA^dsxF.1^, the *dsxF* locus was amplified with the 1114A.S40 (5’ CATATGGTGTTATGCCACGTTCAC 3’) and 1114A.S41 (5’ CGGAAAGTTTATCATCCACTCTGAC 3’) primers, and the amplicon was digested with Acil (NEB cat#R0551S). To identify mutations by gRNA^fle.7^, the *fle* locus was amplified with the 1154A.S23 (5’ CTCAGCAAGCAGTATGCCAAC 3’) and 1154A.S8 (5’ GTTGAACGCTTCGTCGTACG 3’) primers, and the amplicon was digested with BseYI (NEB cat# R0635S). All PCR reactions were performed using Q5 HotStart DNA polymerase (NEB, Cat. No./ID: M0493L). Digestions were performed at 37C for 1 hour, then run on 1% agarose gel at 120V for 25 minutes. Undigested bands corresponding to mutant PCR products were gel extracted with the Zymoclean Gel DNA Recovery Kit (Zymo Research, Cat. No./ID: D4007), then cloned into pJET vectors (Thermo Scientific, Cat. No. / ID: K1231), transformed into chemically competent *E. coli* (Promega, JM109), and plated on LB-Ampicillin plates. Plates were sent for Sanger Colony sequencing with universal primers PJET1-2F (5’ CGACTCACTATAGGGAGAGCGGC 3’) and/or PJET 1-2R (5’ AAGAACATCGATTTTCCATGGCAG 3’), with each colony representing a single PCR amplicon from an individual mutant allele. Sequences were compared to the reference genome sequences of AGAP006241, AGAP008622, AGAP004050, AGAP013051 for *zpg*, β*2-tubulin, dsxF*, and *fle* respectively (**Figure S3**).

### Population suppression assays

On day 1 of experimentation, cages were seeded with 0, 50, 100, 250, or 500 virgin 2-4 day old pgSIT males (for release ratios of 0:1, 1:1, 2:1, 5:1 and 10:1 respectively) intermixed with 50 virgin 2-4 day old wild type males. Then 50 2-4 day old virgin wild type females were then aspirated into the cage. Adults were allowed to mate *ad libitum,* then blood-fed on a mouse on day 6. A wet filter paper (oviposition site) was provided on day 8, and eggs were allowed to develop and hatch undisturbed. Hatched larvae were counted on days 11, 12, and 13 and screened for fluorescence, which would indicate a fertile pgSIT father. Egg shells were counted on days 13 and 14. Only replicates which yielded >1000 eggs were included to guarantee ample representation of male contribution. Each data point represents the counts from a single distinct cross cage; individual cages were not scored multiple times.

### Mathematical modeling

We used the MGDrivE 2 framework (64) to simulate releases of *A. gambiae* pgSIT eggs to suppress mosquitoes in the Upper River region of The Gambia. MGDrivE 2 is a modular framework for simulating releases of genetic control systems in spatially-structured mosquito populations which includes modules for inheritance, life history, and epidemiology. The inheritance pattern of the pgSIT system was modeled within the inheritance module of MGDrivE (101). Based on laboratory data, we assumed the pgSIT system in *A. gambiae* would induce complete male sterility and female inviability, with inviability being manifest at pupation through female androgenization. We assumed that pgSIT eggs would be introduced into the environment in cups with sufficient water volume and larval resources such that larval mortality would be density-independent. Survival of eggs released in cups was determined by expected juvenile life stage durations and daily mortality rates (**Table S8**) leading to a viable emergence rate of 26% for male eggs. Once emerged, offspring of pgSIT sterile males produce offspring with 0% pupatory success, but which are viable at the larval stage, and hence contribute to larval density-dependent mortality. To be conservative with our predictions, we assumed a 25% reduction in both pgSIT male mating competitiveness and lifespan compared to wild-type males, although there is no current laboratory data to suggest a fitness disadvantage (52, 53).

The MGDrivE 2 framework (64) models the development of mosquitoes from egg to larva to pupa to adult with overlapping generations, larval mortality increasing with larval density (102), and a mating structure in which females retain the genetic material of the adult male with whom they mate for the duration of their adult lifespan. Life history of *A. gambiae* was modeled using standard bionomic parameters (**Table S8**) and seasonality in larval carrying capacity driven by rainfall data from the Upper River region of The Gambia (https://www.chc.ucsb.edu/data/chirps). To smooth the seasonal profile of the raw rainfall data, we leveraged a Fourier analysis-based approach that involves fitting a mixture of sinusoids to the raw data (https://github.com/mrc-ide/umbrella). Entomological data from the Upper River region (66) suggested vector breeding sites in this region are substantially more abundant in the rainy season than in the dry season, suggesting larval carrying capacity in the dry season was ~10% that of the peak rainy season. We calibrated the model to malaria prevalence data from a randomized-controlled trial of mass drug intervention in the Upper River region (65) by linking MGDrivE 2 (64) to the Imperial College London (ICL) malaria model (103, 104) by allowing forces of infection (i.e., the probability of infection from mosquito-to-human and human-to-mosquito per individual per unit time) to be exchanged between the two models. Weekly releases of pgSIT *A. gambiae* eggs were simulated from the beginning of the rainy season (June 1st), for a variable number of weeks and release sizes.

### Ethical conduct of research

All animals were handled in accordance with the Guide for the Care and Use of Laboratory Animals as recommended by the National Institutes of Health and approved by the UCSD Institutional Animal Care and Use Committee (IACUC, Animal Use Protocol #S17187) and UCSD Biological Use Authorization (BUA #R2401).

## Data availability

Complete sequence maps and plasmids are deposited at Addgene.org (200640). All Nanopore sequencing data has been deposited to the NCBI sequence read archive (PRJNA978105). All data used to generate figures are provided in the Supplementary Materials/Tables. *A. gambiae* transgenic lines are available upon request to O.S.A.

## Acknowledgments

We thank Judy Ishikawa for helping with mosquito husbandry and Akshay Bharadwaj for laboratory assistance. This work was supported in part by funding from an NIH award (R01AI151004), EPA (Award #RD84020401), and an Open Philanthropy award (309937-0001) awarded to OSA; and by funds from the Bill & Melinda Gates Foundation (INV-017683) awarded to JMM. The views, opinions, and/or findings expressed are those of the authors and should not be interpreted as representing the official views or policies of the U.S. government. Figures were made with Biorender.com

## Author Contributions

O.S.A, A.L.S., R.A.A., and J.J.P conceptualized and designed experiments. R.A.A., J.J.P, A.L.S., M.C., and S.C., performed molecular analyses; R.A.A., J.J.P, A.L.S., M.C., and S.C. performed genetic experiments. A.M., H.M.S.C., and J.M.M. performed mathematical modeling; I.A. performed bioinformatics. All authors contributed to analyzing and compiling the data, and writing and approving the final manuscript.

## Competing Interests

O.S.A is a founder of Agragene, Inc. and Synvect, Inc. with equity interest. The terms of this arrangement have been reviewed and approved by the University of California, San Diego in accordance with its conflict of interest policies. All other authors declare no competing interests.

## SUPPLEMENTARY FIGURES

**Figure S1.**
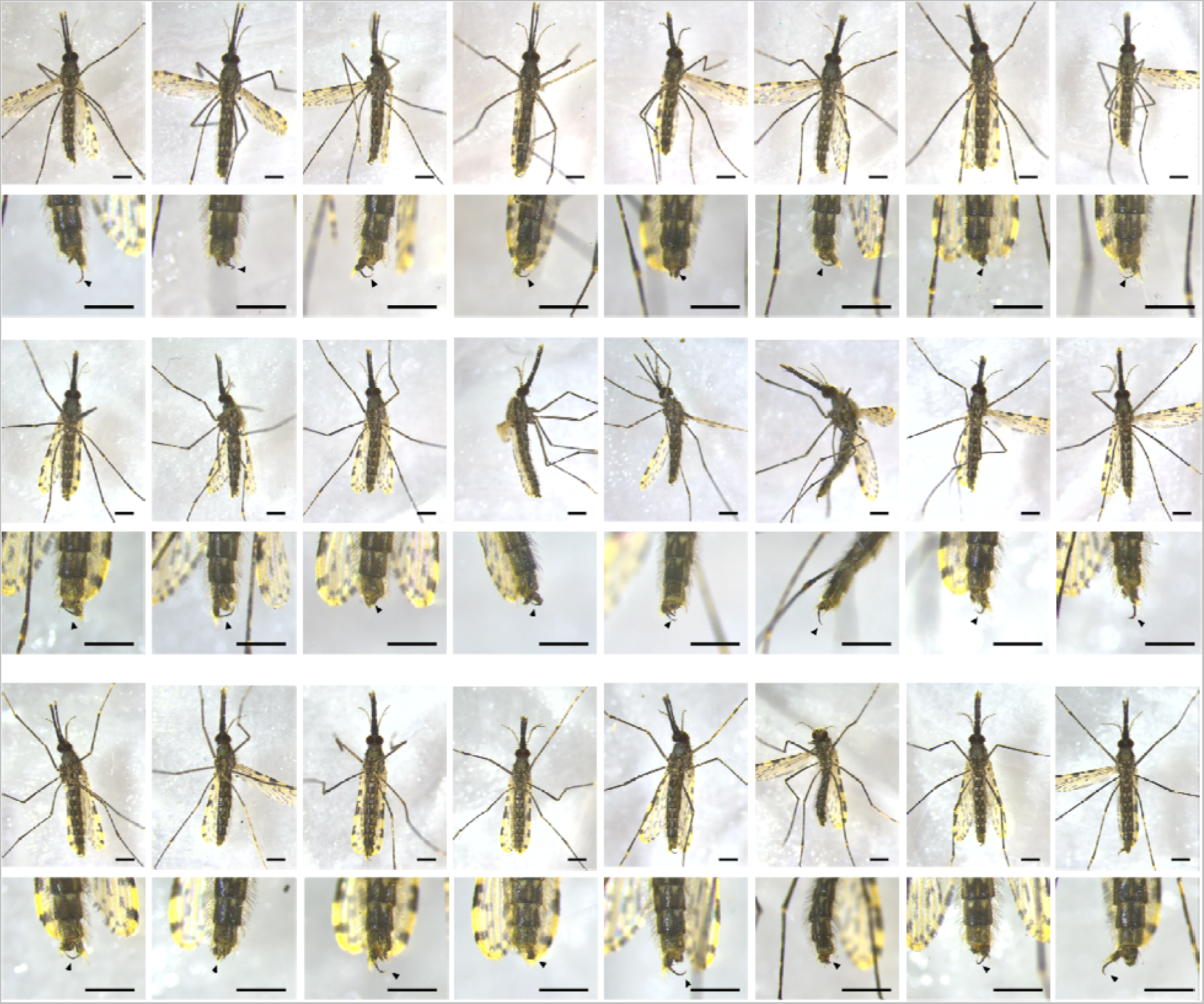
Intersex genitalia of F1 (+/gZBD; +/Cas9) females. (+/gZBD; +/Cas9) hybrid F1 female offspring from gZBD males crossed to Cas9 females. Evidence of *dsxF* mutagenesis as indicated by the presence of development of a male clasper (black arrow) on the female genital appendage. Scale bars indicate 1mm. 24 distinct intersex females shown.

**Figure S2.**
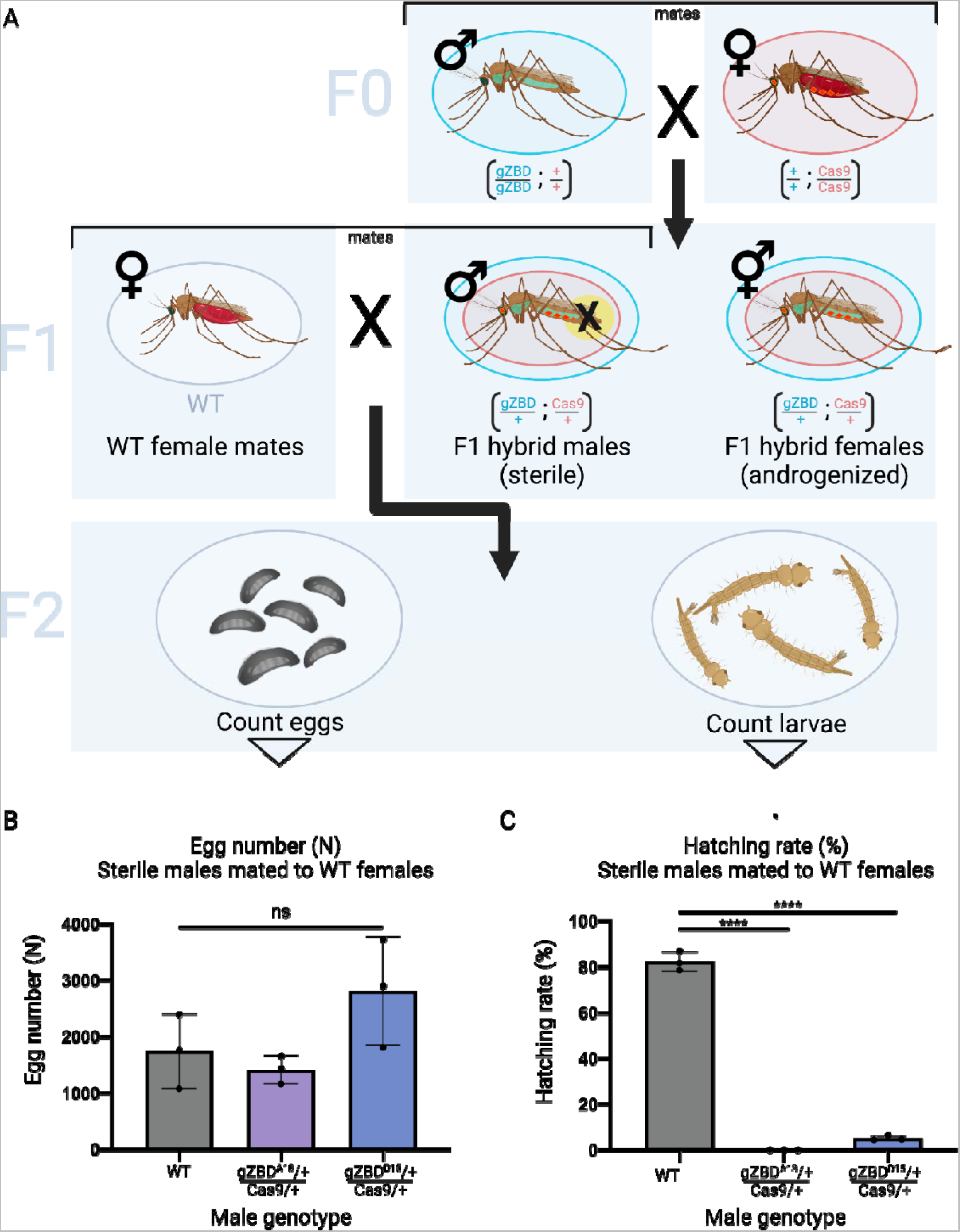
Sterility of (+/gZBD; +/Cas9) F1 males. **A)** To generate F1 males, F0 gZBD males and Cas9 females are crossed together. In the F1 generation, females are expected to be androgenized and males are expected to be sterilized. To test male sterility, F1 (+/gZBD; +/Cas9) males are mated to wild type virgin females. Among the F2 offspring the number of larvae and eggs are counted to measure hatching rate (n% = 1 day old larvae / total number of eggs laid). **B)** Crosses of 50 (+/gZBD^A18^; +/Cas9), (+/gZBD^D15^; +/Cas9), or wild type control males mated to 50 wild type females were performed, and the number of eggs laid is reported. Egg number is nonsignificantly different between all groups (p=0.1057, One Way ANOVA). Mean and SD shown. **C)** The hatching rate of broods from the above described crosses is reported. Hatching rate is significantly lower than wild type controls for both sterile male groups (both p=<0.0001, One Way ANOVA, Dunnett’s multiple comparisons test). Mean and SD shown. Observed larvae were present at the Mendelian ratios expected of transgenic fathers, and the mean hatch rate of these groups is reported in gray.

**Figure S3.**
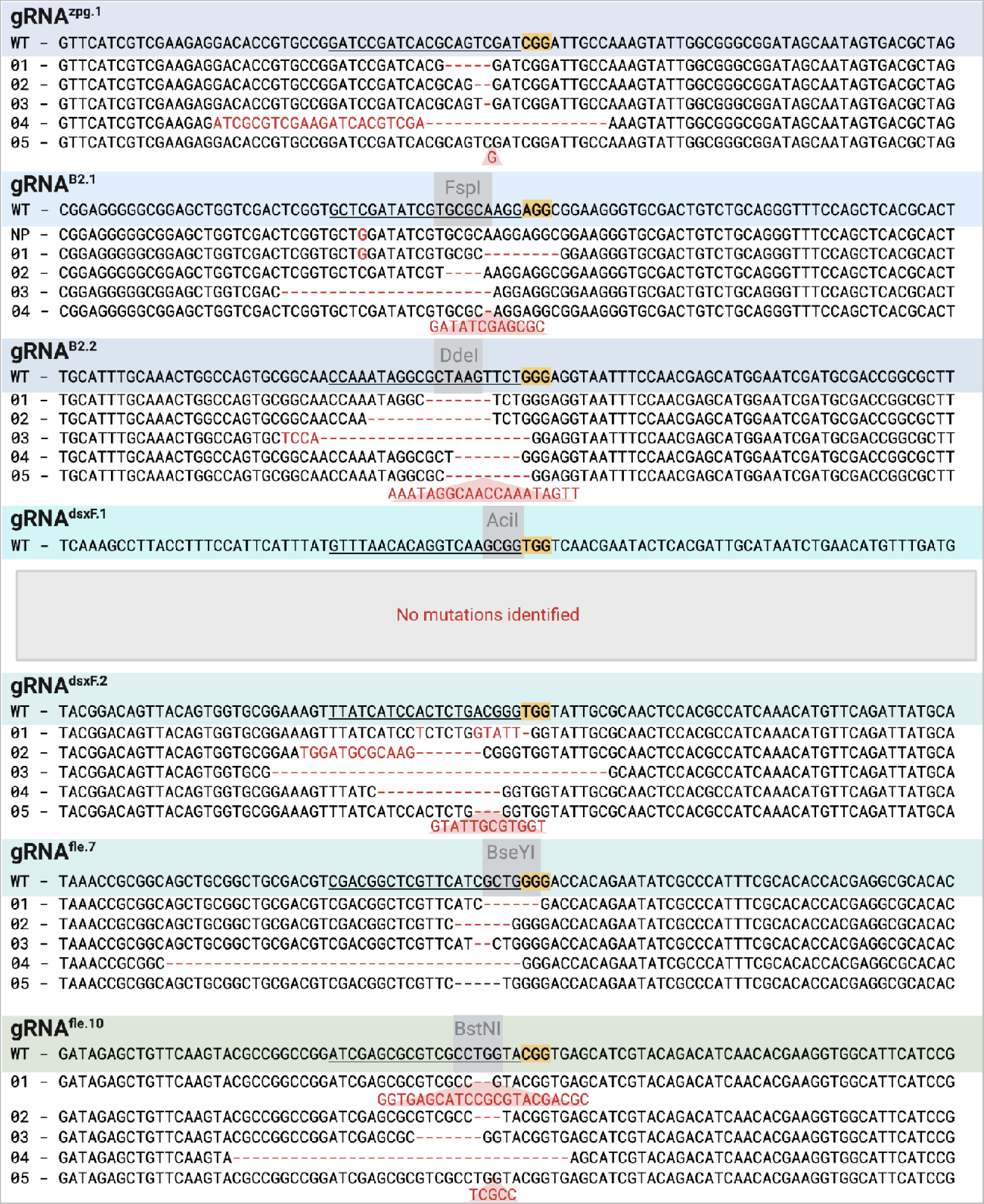
Target site mutations in pgSIT embryos. Each gRNA target site is underlined with the PAM highlighted in yellow. To isolate mutations under some gRNAs, the target site was PCR-amplified and digested with a restriction enzyme whose recognition sequence lies near the CRISPR cut site (grey box). Failure to cleave the PCR product was indicative of CRISPR mutations, and individual amplicon sequencing confirmed the sequence. Mutant reads are derived from (+/gFLE;gZBD^D15^; +/Cas9) individuals are marked with 0#. Naturally occurring polymorphisms are denoted by NP.

**Figure S4.**
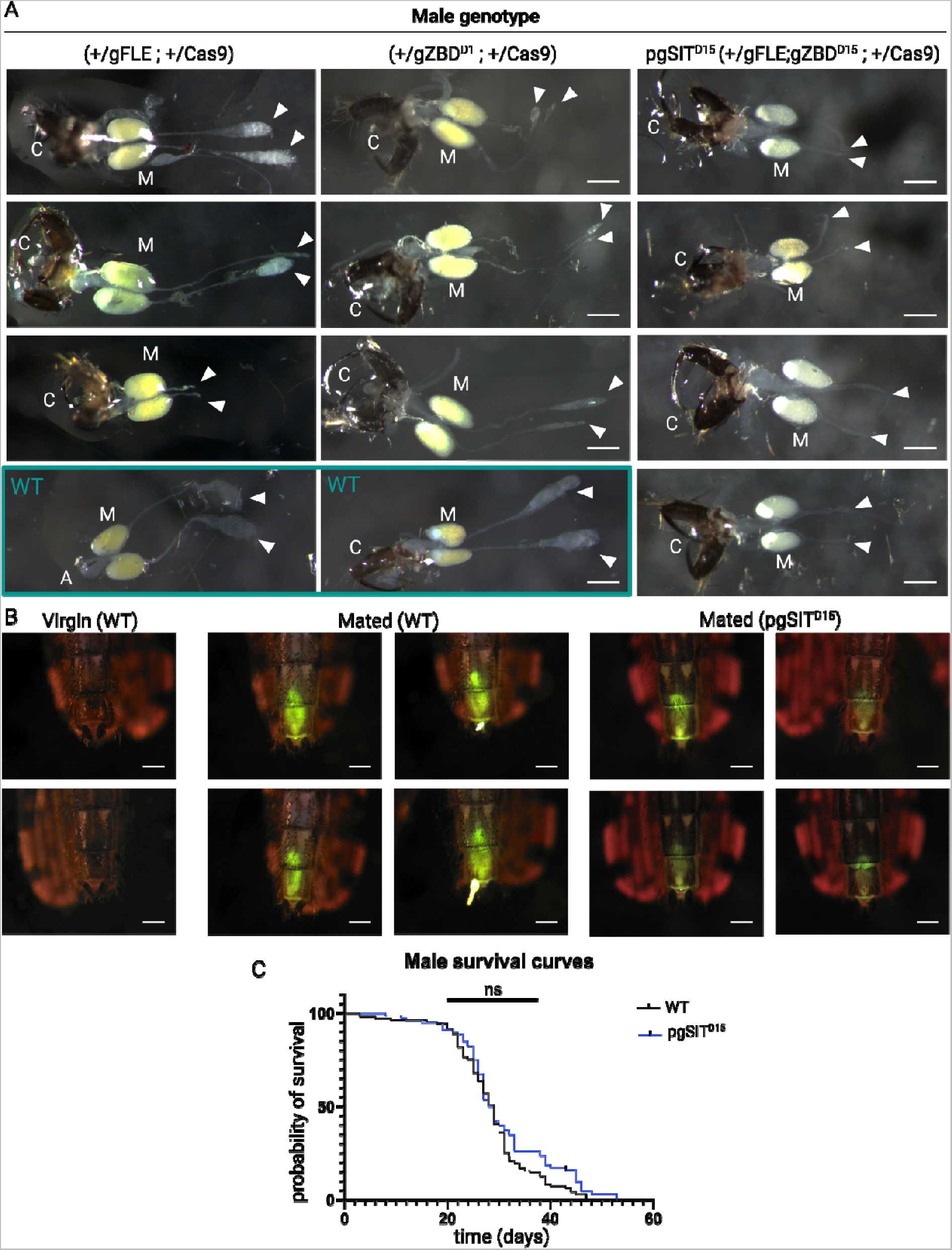
pgSIT^D15^ male reproductive and fitness phenotypes. **A)** Dissections of lower reproductive tracts from three (+/gFLE; +/Cas9), three (+/gZBD^D15^; +/Cas9) and four pgSIT^D15^ (+/gFLE;gZBD^D15^; +/Cas9) males shown, highlighting the presence or absence of testes. In the (+/gFLE; +/Cas9) group, tissues in which both, a single, and neither testes develop are shown. Wild type control reproductive tracts are shown boxed in teal. Both wild type reproductive tracts with attached and removed claspers are shown for reference; a clasper-less image is shown for reference to other published works, and an image with the claspers is included to show the presence of this important reproductive appendage, consistent with the other panels shown. Aedaegus is labeled with an A in the clasper-less sample. Claspers are denoted with C, Male Accessory Glands (MAGs) are denoted with M, the location at the end of the vas deferens where testicular tissue should be developed is marked with an arrow. Lighting brightness, position, white balance, and exposure time were not controlled for, therefore no conclusions regarding the brightness or color of tissues should be made. Scale bars denote 200 µm. **B)** The yellow-green autofluorescent mating plug is visible within the female lower reproductive tract and is sufficiently bright to be observed through the cuticle. In some females, a portion of the plug protrudes externally beyond the gonotreme. Virgin controls are unmated and therefore have no autofluorescent plug. Lighting brightness, position, and exposure time were not controlled for, therefore no conclusions regarding the brightness or color of tissues should be made. Scale bars denote 200 µm. **C)** Male survival curves. Male pgSIT^D15^ and wild type adults were monitored daily for death. pgSIT^D15^ survival differs non-significantly differently from wild type (Log-rank Mantel-Cox).

**Figure S5.**
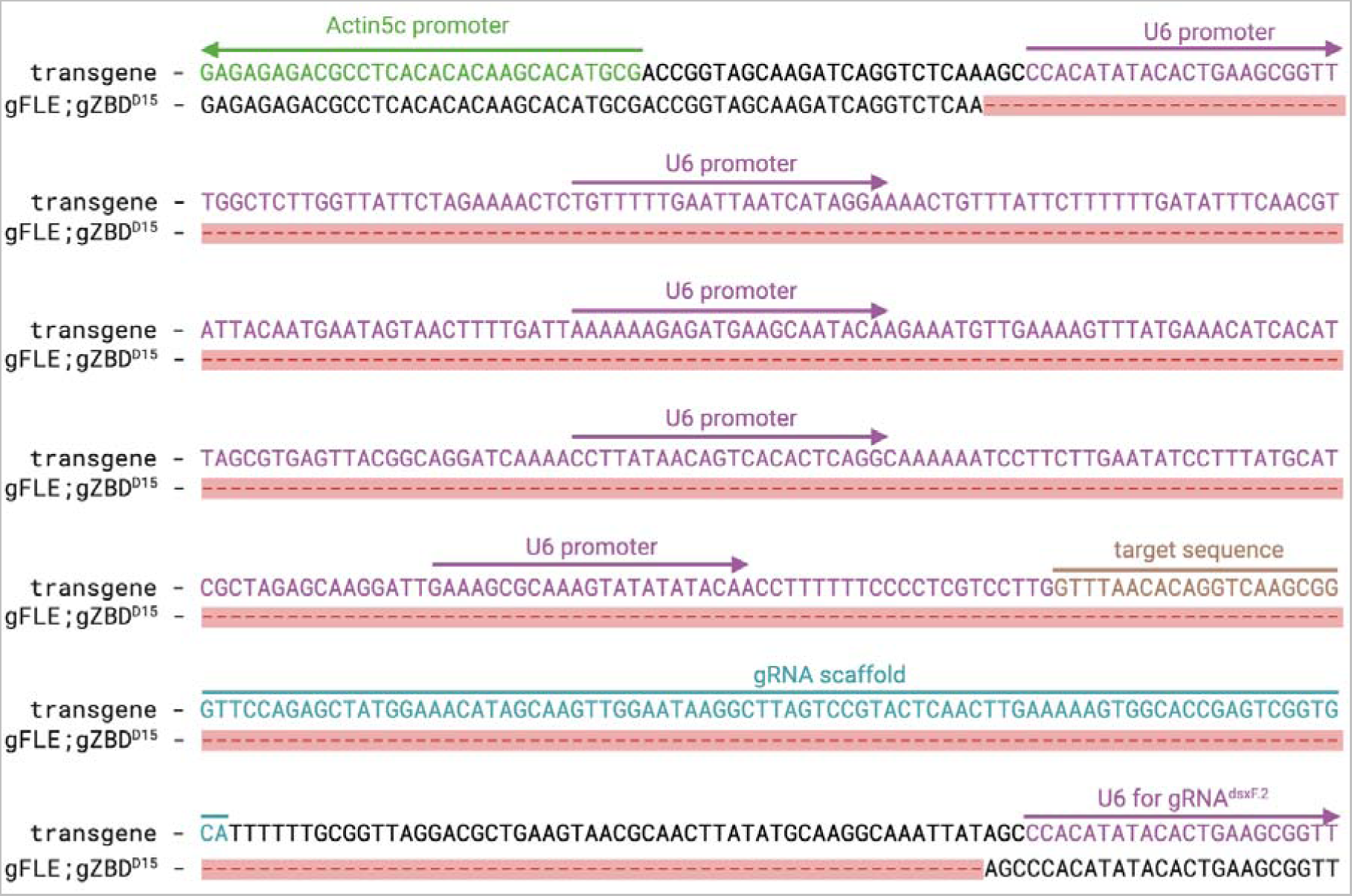
Sequencing of gRNA^dsxF.1^ coding sequence and surrounding region in pgSIT^D15^ individuals. The gZBD^D15^ line showed the same mutation, not shown. The beginning of the adjacent *Act5C* promoter is shown in green, transcribing towards the left in the antisense. The sequence of the *U6* promoter is shown in purple. The sequence corresponding to the gRNA target sequence is shown in gold. The sequence corresponding to the gRNA scaffold is shown in teal. A portion of the beginning of the *U6* promoter which transcribes the adjacent gRNA^dsxF.2^ is also shown. The mutant sequence is shown in black with red dashes denoting the deletion.

## Text S1. Calculation of sort:release ratio

The rate limiting step for many vector control releases is the ability to sort releasable males from undesirable females. With current technology, sorting can be achieved two ways: fluorescently and optically. Fluorescence based sorting relies on a modified cell sorter machine, a COPAS, to sort up to 40,000 freshly hatched nascent larvae per hour (https://malariajournal.biomedcentral.com/articles/10.1186/1475-2875-11-302). In contrast, optically-based sorting relies on computerized visual identification of adult males vs females, and has a significantly slower sort speed (exact number per hour not disclosed)(https://www.nature.com/articles/s42003-023-05030-7 this is verily’s optical sorting papers in aedes). Not only do these technologies vary significantly in their sort speed, but optical sorters necessitate rearing of the undesirable sex to adulthood, wasting significant resources on husbandry.

Most vector control systems rely on direct isolation of males in the generation to be released (with the exception of RIDL systems). This yields a 2:1 sort:release ratio because one male and one female are sorted for every singular male released. This is the most common sorting:release ratio for most vector control systems, and is a ratio which requires a significant number of sorting machines to be able to achieve the scale required for mass releases. However this is in contrast with systems which automatically eliminate the females in the released generation, such as Ifegenia and pgSIT. In such systems, the F0 generation is sex sorted, and the F1 generation is released, improving throughput by orders of magnitude because the fecundity of F0 females is so high. Specifically, in pgSIT, 4 adult mosquitoes must be sorted - Cas9 males and gRNA females each from a pool of both sexes - to yield one fertilized F0 female, who can conservatively produce 400 eggs in her lifetime (though some say upwards of 1,000 eggs (https://targetmalaria.org/wp-content/uploads/2020/11/Ecology_FS_EN_Anopheles-gambiae-s.l.-morphology_August20.pdf). This yields a 4:400, or 1:100 sort:release ratio, and with half of the eggs being male, gives a final sort:release ratio of 1:50.

## SUPPLEMENTARY TABLES

**Table S1:** Quantification of *dsxF* knock-out phenotype in (+/gZBD; +/Cas9) females from gZBD males X Cas9 females

**Table S2:** Raw larvae and egg counts from (+/gZBD; +/Cas9) males X wild type females for male sterility assay (Presented in **Figure 2B**,**C**)

**Table S3**: Raw pupae counts of offspring from homozygous PgSIT X Cas9 in both directions - PgSIT males X Cas9 females results in high lethality in offspring

**Table S4**: Raw pupae counts from homozygous PgSIT female X Cas9 male for female elimination assay (Presented in **Figure 1B**)

**Table S5**: Raw larvae and egg counts from F1 pgSIT males (+/gFLE;gZBD; +/Cas9) X wild type females for Male sterility assay (Presented in **Figure 1C**,**D**)

**Table S6**: Raw counts of dead F1 pgSIT adult males (+/gFLE;gZBD; +/Cas9) for Male adult survival assay (Presented in **Figure S4C**)

**Table S7**: Raw larvae and egg counts from F1 pgSIT males X wild type males X wild type females (at different ratios) for Population Suppression Assays (Presented in **Figure 2B,C**)

**Table S8**: Model parameters describing pgSIT construct, mosquito bionomics and malaria epidemiology for simulated releases in Upper River region, The Gambia.

